# Impacts of climate change on California’s rangeland production: sensitivity and future projection

**DOI:** 10.1101/2022.02.06.479265

**Authors:** Han Liu, Yufang Jin, Leslie M. Roche, Anthony T. O’Geen, Randy A. Dahlgren

## Abstract

Rangelands support many important ecosystem services and are highly sensitive to climate change. Understanding temporal dynamics in rangeland gross primary production (GPP) and how it may change under projected climate change, including more frequent and severe droughts, is critical for ranching communities to cope with future changes. Covering ~10% of California’s climatologically and topographically diverse landscapes, annual rangelands express varying sensitivity to precipitation fluctuation and warming. Herein, we examined how climate regulates temporal dynamics of annual GPP in California’s annual rangeland across scales, based on 20 years of satellite record derived GPP at 500-meter resolution since 2001. We built gradient boosted regression tree models for 23 ecoregion subsections in our study area, relating annual GPP with 30 climatic variables and disentangling the partial dependence of GPP on each climate variable. Our analysis showed that GPP was most sensitive to growing season precipitation amount; GPP decrease as much as 200 g C/m^2^/yr when growing season precipitation decreased from 400 to 100 mm/yr in one of the driest subsections. We also found that years with more evenly distributed growing season precipitation had higher GPP. Warmer winter minimum air temperature enhanced GPP in approximately two-thirds of the subsections. In contrast, average growing season mean and maximum air temperatures showed a negative relationship with annual GPP. When forced by downscaled future climate projections, changes in future rangeland productivity at the ecoregion subsection scale were more remarkable than at the state level; this suggests rangeland productivity responses to climate change will be highly variable at the local level. Further, we found large uncertainty in precipitation projections among the four climate models used in this study. Specifically, drier models predicted a larger degree of reduction in GPP, especially in drier subsections. Our machine learning-based analysis highlights key regional differences in GPP vulnerability to climate and provides insights into the intertwining and potentially counteracting effects of seasonal temperature and precipitation regimes. This work demonstrates the potential of using remote sensing to enhance field-based rangeland monitoring and, combined with machine learning, to inform adaptive management and conservation within the context of weather extremes and climate change.

## 1. Background

Gross primary production (GPP) is a widely used metric for quantifying photosynthesis rate at the ecosystem scale. Recent studies have shown increasing interannual variability in GPP globally, with 57% of the variability contributed by rangeland-dominated ecosystems (Zhang et al., 2016). In semi-arid rangelands, precipitation is, in most cases, the primary controller of GPP fluctuations (Zhang et al., 2016). With projected increasing precipitation variability, the interannual variability of GPP is also predicted to increase (Zscheischler et al., 2014). Changes in rangeland GPP may impact many important ecosystem services, including carbon storage, water resource protection, biodiversity, and wildlife habitat (Li, 2012; Roche et al., 2015). Therefore, monitoring GPP and understanding its variability may be a useful benchmark indicator of ecosystem response to climate change Rangelands cover more than 30% of the global land area and are biologically diverse landscapes that include grasslands, shrublands, woodlands, wetlands, and deserts (Gyde Lund, 2007). In California, annual rangelands, which include open grasslands and woodlands dominated by an understory of herbaceous annual plants, are the primary forage source for the State’s $3 billion livestock industry (California Department of Food & Agriculture, 2015). These annual rangelands comprise approximately 10 million acres, encompassing portions of the Central Valley, the Coast Range and Sierra Foothill Region (FRAP, 2017). California rangeland are highly variable, in part, because the plant community phenology, consisting of annual grasses and forbs, is dynamically linked to seasonal weather patterns (Stromberg, Corbin & D’Antonio, 2007). For example, the growing season is initialized by fall rains (September-December) exceeding 1.25-2.5 cm during a seven-day period, which promotes germination. Over winter (December-February), plant growth slows due to lower air temperatures and lower solar radiation. In late winter/early spring (February-April), rapid growth begins as air temperature increases (>7.2°C) and solar radiation intensifies. Finally, in mid-late spring (April-June), plant production reaches peak biomass once root zone moisture is depleted. Thus, California rangeland GPP may vary significantly seasonally, from year-to-year, and across locations (Liu et al., 2021) with respect to short-term changes in precipitation, air temperature, and solar radiation.

Over the past decades, researchers have relied on regression and/or correlation analyses to examine drivers regulating variability in annual GPP and plant biomass production on rangelands. Taking advantage of the high correlation between absorbed photosynthetically active radiation (APAR) and GPP, Jin and Goulden (2014) reported GPP was positively related to annual precipitation, with higher sensitivity in grasslands than woody shrublands in California. A few other studies also reported positive relationships between rangeland GPP and annual precipitation amount in Northern China (Guo et al., 2015), North America (Wu & Chen, 2012), and across the entire globe (Huang et al., 2016). However, annual GPP is affected by more than precipitation amount. Seasonal distribution of annual precipitation events also has a significant impact on rangeland GPP by regulating the active growing season (Xu & Baldocchi, 2004). Dass, Rawlins, and Kimball (2016) found that while a warming air temperature is the primary reason for recent GPP increases in northern Eurasia’s shrublands, they also found warming air temperature drove decreased GPP in the northern Eurasia’s southwestern grasslands. In California, rangeland production is predicted to increase as air temperatures become warmer in the San Francisco Bay Area as early as mid-century (Chaplin-Kramer & George, 2013); however, another study predicted a 14 to 58% decrease in rangeland production across California using a precipitation-based model (Shaw et al., 2011). Previous studies showed variations in GPP were related to the combined effects of various climatic variables, and the relationship between GPP and climatic variables may vary across different locations. Thus, to fully understand variations in GPP and predict future trends, it is necessary to build individual models with respect to different regions and consider a suite of relevant climate variables, including precipitation amount, seasonal distribution, air temperature, and solar radiation.

MODIS has been collecting terrestrial observations for more than twenty years on a daily basis since the launch of Terra satellite in 1999 (Savtchenko et al., 2004). GPP products derived from MODIS satellite images at 1-km spatial resolution are available at an eight-day interval for the entire globe. This twenty-year record allows for a more robust analysis of interannual variability in GPP. Machine learning models have also become more interpretable while keeping the powerful non-linear multi-parameter model fitting functionalities (Molnar, 2019). Advances in both remote sensing technologies and data analysis are providing novel, cost-effective opportunities to obtain data at management-relevant scales for terrestrial-ecosystem research, which has otherwise relied on traditional statistical methods and field-based monitoring that require substantial time and labor resources (Briske et al., 2017). These advances are key because traditional statistical models lack the ability to disentangle non-linear responses and interactions among variables.

In this study, we used 20 years of remote sensing records and machine learning methods to improve our understanding of temporal variability in rangeland annual GPP and how its sensitivity to changes in weather and climate may vary across different ecoregions in California. Specifically, we addressed the following questions: 1) What climatic variables are the primary drivers for interannual GPP variation, and how sensitive is GPP to each variable? 2) To what extent will rangeland GPP change by mid-century and end-of-century across California? and 3) How do changes in each climate variable contribute to future changes in GPP?

## 2. Data & Methods

### 2.1 Study area

Our study focused on California annual rangelands in four major ecoregions, based on ECOMAP (Cleland et al., 2007): Northern California Coast Ranges (NCCR), Northern California Interior Coast Ranges (NCICR), Central Coast Ranges (CCR), and Sierra Nevada Foothills (SNF) (Fig. 1a). This area spans from 33° to 41° N and 118° to 124°, with a wide range of microclimates because of their complex geographic diversity. These annual rangelands feature a Mediterranean climate with hot, dry summers and mild, moderately wet winters. The active growing season of annual rangelands coincides with the rainy season: germination typically happens in late fall (November), and senescence occurs quickly after the rainy season ceases in late Spring or early Summer (April-May).

**Figure 1.**
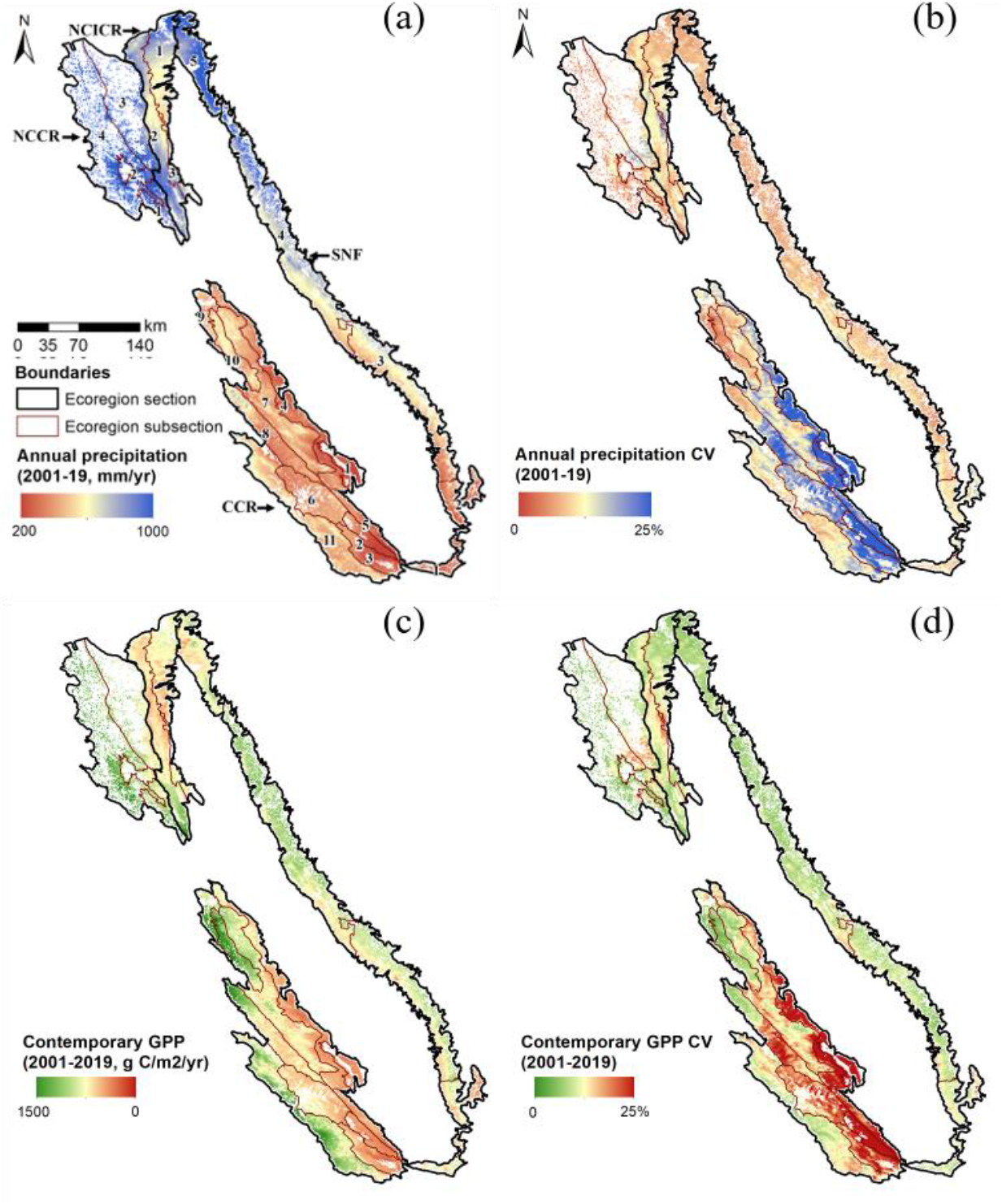
Statewide distribution of long-term-mean (a) precipitation and (b) GPP during 2001-2019 over California’s annual rangelands. Maps of year-to-year variability, as quantified by the coefficient of variation (CV), are also shown (c, d). The study area encompasses four ecoregions (black lines) that are further divided into 23 subsections (orange lines).

Precipitation varies considerably across regions, with its long-term-mean ranging from 160 mm/yr in the CCR to 2290 mm/yr in the NCCR (Fig.1a). Coastal regions are relatively cooler in the summer and receive higher amounts of precipitation than inland regions, especially those in rain shadows. Across the study area, air temperature increased from the coast to inland, north to south, and from valleys to mountains. Based on topographic variables, soil types, and climatic conditions, the four major ecoregions are further divided into 23 ecoregion subsections (Fig. 1a) (Cleland et al., 2007).

### 2.2 Satellite and climate data

We obtained the version 6 gap-filled GPP product (MYD17A2HGF) at 500-m resolution every eight days from Moderate Resolution Imaging Spectroradiometer (MODIS) Terra and Aqua during 2000-2019 (https://earthdata.nasa.gov/). MODIS provides frequent multispectral moderate-resolution (250-500 m) observations, revisiting the globe once to twice per day since 2000. We calculated water year (October-September) cumulative GPP from the 8-day product. A long-term mean annual GPP layer was derived from 2001-2019 as a measure of spatial variation in forage production. The annual cumulative layers were further resampled to 1-km resolution with a linear method to match Daymet grids.

Meteorological data during 1986-2019 came from Daymet hosted at the Oak Ridge National Laboratory Distributed Active Archive Center (Thornton et al., 2016). The gridded daily surface weather parameters at a 1-km spatial resolution were interpolated from more than 8,000 meteorological stations, based on a digital elevation model (Thornton et al., 2016). We aggregated daily precipitation, min/max/mean air temperatures, and solar radiation to monthly and seasonal averages, as detailed in Table 1. To quantify seasonal precipitation variation between the driest and wettest month during the growing season (November to May), a Precipitation Concentration Index (PCI) was calculated as the ratio of the sum of squares of the wet season monthly precipitation (November to May) and squared water year precipitation times a scaling factor 58.0 (Oliver, 1980; Sloat et al., 2018).

**Table 1.** List of variables used as predictors for machine learning GPP modeling. Seasonal variables include averages during growing season (GS, October-May), early, mid- and late seasons (November-January, January-March, March/April-June), Winter refers to December to February, which was further split into early and late Winter (November-January and January-March).

| Variable type (unit) | Variable names and abbreviations | Data source/reference |
| --- | --- | --- |
| Precipitation amount (mm) | Seasonal cumulative precipitation: GS_ppt, EarlyS_ppt, MidS_ppt, LateSApr_ppt, LateSMar_ppt<br>Monthly cumulative precipitation: M11_ppt, M12_ppt, M01_ppt, M02_ppt, M03_ppt, M04_ppt, | Daymet (Thornton et al., 2016) |
| Precipitation distribution (unitless) | Precipitation concentration index during wet season (Nov.-May): PCI | Daymet (Thornton et al., 2016; Oliver, 1980) |
| Minimum air temperature (°C) | Growing season and monthly average minimum air temperature: GS_tmin, M11_tmin, M12_tmin, M01_tmin, M02_tmin, M03_tmin | Daymet (Thornton et al., 2016) |
| Mean air temperature (°C) | Seasonal average mean air temperature: GS_tave, Winter_tave, WinterEarly_tave, WinterLate_tave<br>Monthly average mean air temperature: M11_tave, M12_tave, M01_tave, M02_tave, M03_tave, | Daymet (Thornton et al., 2016)<sup>9</sup> |
| Maximum air temperature (°C) | Growing season average maximum air temperature: GS_tmax | Daymet (Thornton et al., 2016) |
| Incoming solar radiation ($\text{W/m}^2$) | Mean incoming solar radiation during the growing season and during Nov.-Mar.: GS_meansrad, GS_meansrad1103 | Daymet (Thornton et al., 2016) |

For future climate, we obtained daily climate projections downscaled for California from Cal-Adapt Data Server (http://albers.cnr.berkeley.edu/data/scripps/loca/). This dataset is bias-corrected and downscaled to a resolution of 1/16° (~6 km) from 32 global climate models (GCMs) using the Localized Constructed Analogues statistical method (Pierce, Kalansky & Cayan, 2018). The data cover 1950-2005 for the historical period and 2006-2099 for future climate projections under two emission scenarios. In this study, we used projections from four GCMs, that have been reported to have good performance in reproducing California’s historical climate (Pierce, Kalansky & Cayan, 2018): HadGEM2-ES, MIROC5, CNRM-CM5, and CanESM2. Representative Concentration Pathway (RCP) 4.5 (Thomson et al., 2011) is described by the IPCC as an intermediate scenario where emissions peak around 2040 and then decline. A worst-case scenario, RCP 8.5, represents rising emissions throughout the 21^st^ century (Thorne et al., 2017). For each model and scenario projection, we acquired daily mean precipitation, air temperature (minimum, maximum, and mean), and solar radiation layers. A bias adjustment was then applied to the GCM data to match the contemporary record, based on the overlapping Daymet Data and GCM historical simulations during 1986-2005 (20 years). Specifically, we first calculated differences between Daymet and GCM data derived variables (see Table 1) for each year during 1986-2005. Then, we calculated average offsets by taking the mean of these differences. Bias-adjusted GCM data were derived by applying a calculated long-term-mean offset to the original simulations.

### 2.3 Machine learning GPP models

We first examined if there was any significant trend during 2001-2019 in annual rangeland GPP and climatic variables, using the Hamed and Rao Modified Mann-Kendall (MK) trend test (Hamed & Ramachandra Rao, 1998). The trend test was conducted at both ecoregion and 1-km pixel levels, and a significant trend was identified when the corresponding p-value was less than 0.05.

We built machine learning models for each ecoregion subsection to investigate the complex associations between the interannual variability in annual rangeland GPP and climatic drivers. The Gradient Boosted Decision Trees (GBDT) approach was chosen due to its robustness over single learner methods (Friedman, 2001). GBRT is an ensemble method that fits regression trees sequentially, with each tree aimed at minimizing prediction residuals of its predecessors. For each ecoregion subsection, a GBDT model was trained with 70% randomly selected 1-km pixels, using key variables selected from Table 1. The long-term-mean (LTM) GPP at 1-km was also included as an additional independent variable in order to help account for spatial variability in GPP caused by topography and soils.

Feature selection was conducted to remove redundant climatic variables and reduce potential overfitting issues. We first iteratively eliminated variables with respect to their correlation to other variables (i.e., when two variables are highly correlated (r>0.8), removing the variable having the lower distance correlation (Székely, Rizzo & Bakirov, 2007) with annual rangeland GPP. We further removed variables that had low predictive power using the Recursive Feature Elimination with Cross-Validation (RFECV) technique (Guyon et al., 2002). Finally, a permutation importance-based feature selection was performed to further reduce the number of variables. Permutation importance is computed as a decrease in the model performance (for example, measured by R^2^ or RMSE) when a variable is permuted. We sequentially permuted variables with the least permutation importance score and fitted models with the most important three to nine variables. For the purpose of balancing model accuracy and complexity, we kept the top seven variables to build ecoregion subsection-specific machine learning models (Fig. S1).

We investigated how each of the key climatic variables affects interannual rangeland GPP variation using partial dependence plots (PDPs) from the GBDT models. Partial dependence marginalizes the machine learning model over the distribution of other independent variables so that the remaining function describes the relationship between the targeting variable and the dependent variable (Molnar, 2019). The y-axis of PDPs represents the difference between the marginalized prediction and mean GPP. For example, a wide range on the y-axis indicates strong sensitivity of GPP to the target predictor when other confounding factors are excluded.

### 2.4 Future GPP projection

We applied machine learning models, built specifically for each ecoregion-subsection, to the Daymet climate data spanning 2001-2019 and projected climate data from 2040-2099. The results were summarized by taking the means over three-time windows: contemporary (2001-2019), mid-century (2041-2059), and end of century (2081-2099), for the RCP 4.5 and RCP 8.5 emission scenarios.

We quantified changes in GPP by mid-century and end of the century caused by each type of climatic variable (precipitation amount, precipitation distribution, minimum air temperature, mean air temperature, maximum air temperature, and solar radiation). For each type of variable, the attribution was performed by comparing the difference between GPPs predicted for the contemporary period and GPPs predicted with future climate simulation for the target type of variables while keeping all other variables as the contemporary climate. Differences between this newly predicted GPP and the original contemporary GPP indicate the extent to which changes in GPP are attributed to changes in precipitation, air temperature, and solar radiation.

## 3. Results

### 3.1 Temporal patterns of contemporary GPP and climate

Overall, no significant trend in annual GPP was found for the vast majority (approximately 92%) of the study area (Figure 2 and Figure S3a). This is consistent with the absence of a significant trend for precipitation, both in terms of growing season (GS) precipitation amount and monthly distribution. A further examination of those areas that had a significant annual GPP trend indicated that 6% of the study area showed a significant decreasing GPP trend (75±37 g C/m^2^/yr), scattered across all four ecoregions, while only 2% showed an increasing trend, clustered in the northern SNF ecoregion (65±30 g C/m^2^/yr) (Fig. S3a). Unlike precipitation, GS minimum air temperature (T_min_) and maximum air temperature (T_max_) increased in 18% and 10% of the study area, respectively, from 2001 to 2019. While GS T_max_ showed a significant increasing trend in only the CCR, GS T_min_ showed increasing trends in CCR, southern SNF, northern NCICR, and southern NCCR. GS solar radiation showed a decreasing trend in southern NCCR and central CCR, accounting for 2% of the study area (Fig. S3b).

**Figure 2.**
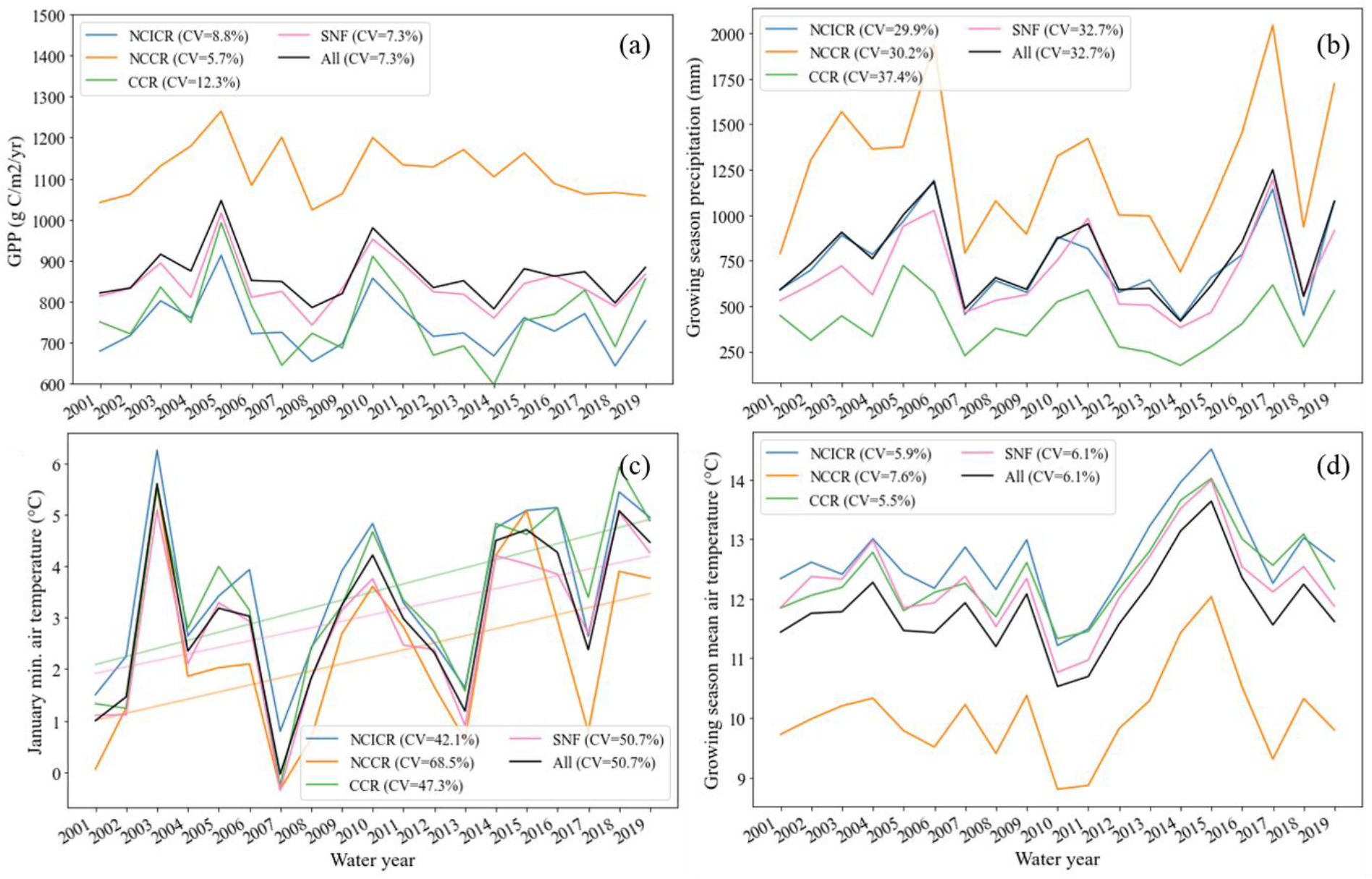
Time series of contemporary (2001-2019) (a) GPP, (b) October-May precipitation (growing season), (c) January T_min_, and (d) October-May T_mean_. The coefficient of variation (CV) is a measure of interannual variability and is calculated as the ratio of standard deviation to mean. Trend lines are shown only for variables with a statistically significant (p<0.05) trend.

MODIS GPP during 2001-2019 water years (820±283 g C/m^2^/yr) varied significantly from year to year, with a CV of 7.3% (Figs. 1d & 2a). Highest productivity was found in 2005 (1017±288 g C /m^2^/yr) and 2010 (946±287 g C /m^2^/yr) (Figure 2a). In contrast, rangeland productivity in water years 2008 and 2014 were much lower, below 750 g C /m^2^/yr. Precipitation fluctuated from year to year with CVs ranging from 30% in NCCR to 37% in CCR (Fig. 1b), and from 397 mm in water year 2014 to 1158 mm in water year 2017 statewide (Fig. 2b). Larger interannual variation in January minimum temperatures was found across all ecoregions (Fig. 2c), although air temperatures during October-May varied to a much lower degree (Fig. 2d).

At the ecoregion subsection scale, interannual variability of GPP was highly correlated with growing season precipitation (e.g., in CCR, NCICR, and southern SNF) (Fig. 3a). Growing season PCI, T_mean_, T_max_, and solar radiation had low to high degrees of negative correlations with GPP across years; the magnitude of these negative correlations also varied spatially across the study area (Fig. 3b & d-f). Furthermore, January T_min_ showed a moderate positive correlation with annual GPP in central CCR (clustered around CCR-6) and in a small area in SNF (mostly in SNF-4) (Fig. 3c).

**Figure 3.**
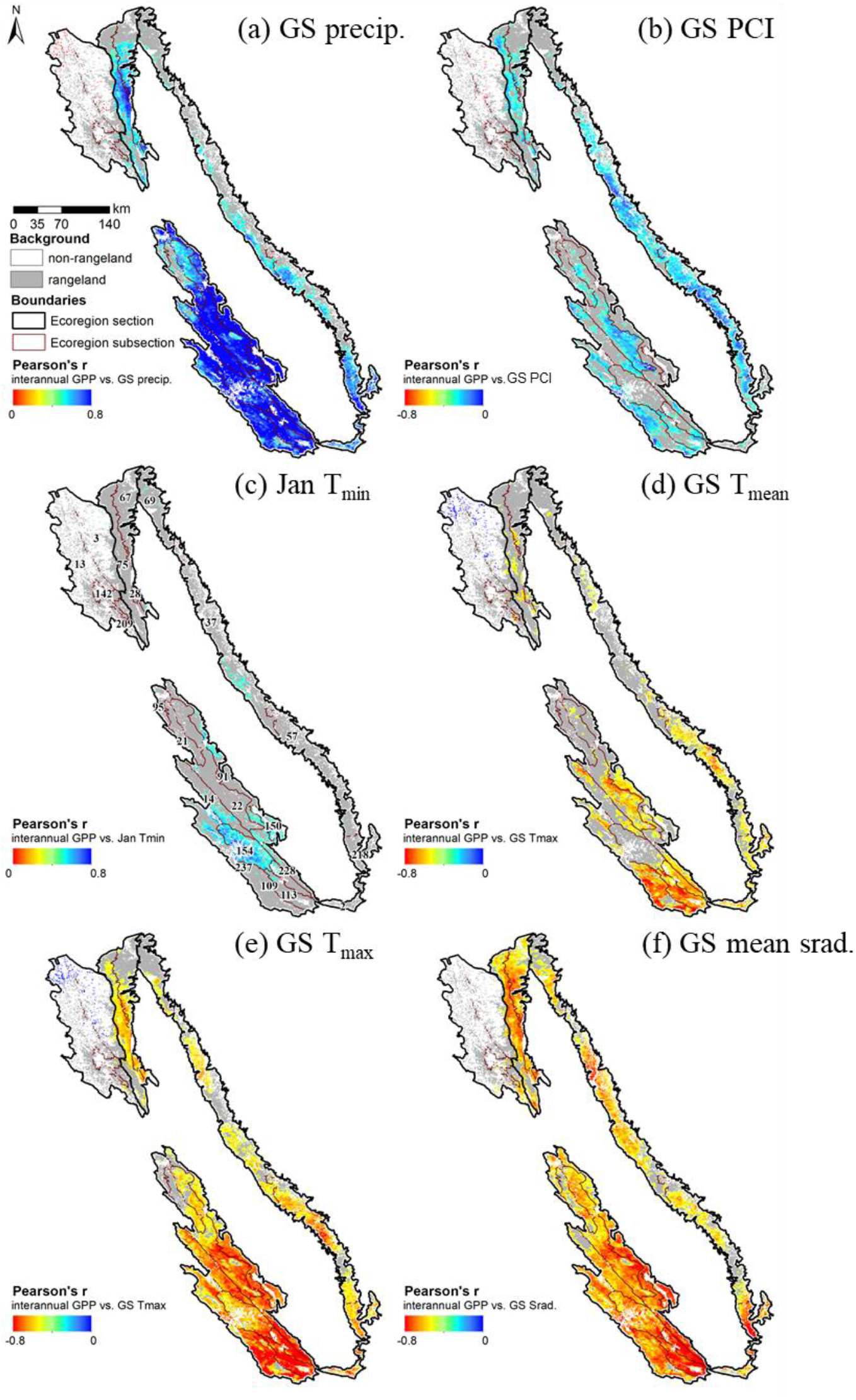
Pearson’s correlation (r) between contemporary GPP and key climatic variables during 2001-2019. Only statistically significant (p<0.05) correlations are shown.

### 3.2 Sensitivity of interannual rangeland GPP to climate

Different climate variables (see Table 2) were selected to model annual rangeland productivity for each of the 23 subsections’ specific GBRT productivity models, using our feature selection method detailed in section 2.3

**Table 2.** Selected variables for each subsection-specific gradient boosted decision trees model after feature selection. See Table 1 for variable abbreviation definitions.

| Subsection | Variables |
| --- | --- |
| CCR-1 | 'GS_ppt', 'MYMGPP', 'M01_tmin', 'M12_tmin', 'M02_tmin', 'M11_tmin', 'M03_tave' |
| CCR-2 | 'GS_tmax', 'MYMGPP', 'GS_ppt', 'M01_tmin', 'M11_ppt', 'M02_tave', 'GS_meansrad' |
| CCR-3 | 'MYMGPP', 'GS_ppt', 'GS_meansrad', 'M01_tmin', 'PCI_wet', 'GS_tmax', 'M11_ppt' |
| CCR-4 | 'MYMGPP', 'GS_ppt', 'M12_tmin', 'M01_tmin', 'M02_tmin', 'PCI_wet', 'M11_tmin' |
| CCR-5 | 'EarlyS_ppt', 'MYMGPP', 'M01_tmin', 'M02_tave', 'M02_ppt', 'LateSMar_ppt', 'PCI_wet' |
| CCR-6 | 'MYMGPP', 'EarlyS_ppt', 'M01_tmin', 'M02_tmin', 'M02_ppt', 'PCI_wet', 'M12_tave' |
| CCR-7 | 'MYMGPP', 'LateSApr_ppt', 'M12_ppt', 'PCI_wet', 'M01_tmin', 'M11_ppt', 'M11_tmin' |
| CCR-8 | 'MYMGPP', 'LateSApr_ppt', 'PCI_wet', 'M01_tmin', 'M11_ppt', 'M12_tmin', 'M12_ppt' |
| CCR-9 | 'MYMGPP', 'PCI_wet', 'LateSApr_ppt', 'EarlyS_ppt', 'M01_tmin', 'M04_ppt', 'M03_tave' |
| CCR-10 | 'MYMGPP', 'PCI_wet', 'M12_ppt', 'LateSApr_ppt', 'M12_tmin', 'M02_tmin', 'M11_ppt' |
| CCR-11 | 'MYMGPP', 'GS_ppt', 'PCI_wet', 'M04_ppt', 'M01_tmin', 'M02_tmin', 'LateSMar_ppt' |
| NCCR-1 | 'MYMGPP', 'M02_tmin', 'M12_ppt', 'PCI_wet', 'M04_ppt', 'M11_ppt', 'M02_ppt' |
| NCCR-2 | 'MYMGPP', 'LateSApr_ppt', 'PCI_wet', 'MidS_ppt', 'M04_ppt', 'M02_ppt', 'M11_tmin' |
| NCCR-3 | 'MYMGPP', 'M02_ppt', 'PCI_wet', 'M04_ppt', 'M01_ppt', 'MidS_ppt', 'M11_tave' |
| NCCR-4 | 'MYMGPP', 'M02_ppt', 'M01_ppt', 'PCI_wet', 'M02_tmin', 'M11_tave', 'M01_tmin' |
| NCICR-1 | 'MYMGPP', 'PCI_wet', 'M11_ppt', 'M12_ppt', 'GS_meansrad', 'M02_tmin', 'GS_tmax' |
| NCICR-2 | 'MYMGPP', 'PCI_wet', 'M12_ppt', 'M11_tmin', 'M02_tmin', 'GS_ppt', 'M04_ppt' |
| NCICR-3 | 'MYMGPP', 'GS_ppt', 'PCI_wet', 'GS_tmax', 'M02_tmin', 'EarlyS_ppt', 'M01_tmin' |
| SNF-1 | 'MYMGPP', 'M12_ppt', 'GS_ppt', 'PCI_wet', 'M04_ppt', 'LateSMar_ppt', 'M01_ppt' |
| SNF-2 | 'MYMGPP', 'LateSApr_ppt', 'PCI_wet', 'LateSMar_ppt', 'M02_ppt', 'M11_ppt', 'M12_ppt' |
| SNF-3 | 'MYMGPP', 'PCI_wet', 'M12_ppt', 'M11_ppt', 'LateSMar_ppt', 'M11_tave', 'LateSApr_ppt' |
| SNF-4 | 'MYMGPP', 'PCI_wet', 'M11_tave', 'M11_ppt', 'LateSMar_ppt', 'M01_tmin', 'M12_ppt' |
| SNF-5 | 'MYMGPP', 'PCI_wet', 'M11_ppt', 'M01_tmin', 'M04_ppt', 'M01_ppt', 'M12_ppt' |

Overall, precipitation-related variables were chosen by more models than other types of variables (Fig. 4b). For example, at least one variable representing precipitation amount was selected for all subsections, either for individual winter and/or spring months or seasonal precipitation, depending on the region. PCI was critical for capturing rangeland productivity variation in 21 out of the 23 subsections, except for the two driest subsections (CCR-1 and CCR-2). Winter minimum air temperature was the third most selected variable (Fig. 4b, in 19 subsections), followed by mean air temperature, which was selected in 14 subsections (Fig. 4a).

**Figure 4.**
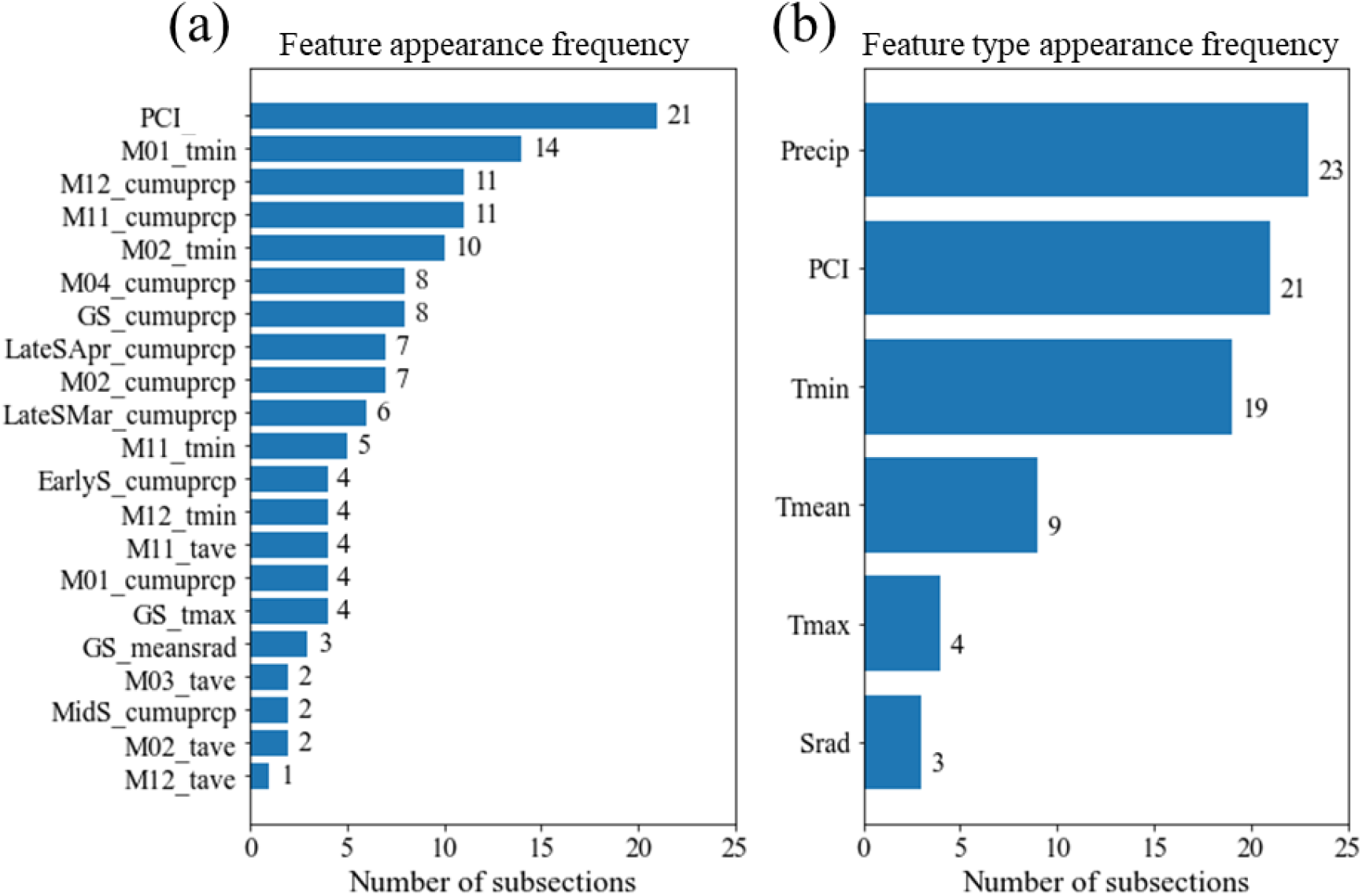
Prevalence of selected (a) climate variables and (b) climate variable groups, summarized among the 23-subsection specific models. For example, an appearance in 21 subsections for precipitation concentration index means PCI was selected as one of the seven variables in 21 subsection models. See Table 1 for variable abbreviation definitions.

Solar radiation was selected in subsections SNF-1, CCR-2 and CCR-3 (Table 2), the most northern and southern subsections of the study area (Fig. 1a). Despite the large latitudinal difference, all subsections had higher annual GPP in years with a lower amount of solar radiation (Fig. 5e).

Predicted GPP was in good agreement with the MODIS GPP product when tested against the validation dataset, with an R^2^ of 0.97 and RMSE of 30.7 g C/m^2^/yr. Mean absolute percentage error (MAPE; calculated by taking the average of percent errors for the absolute value of all observations in the validation dataset) was 4.1%, averaged for the 23 ecoregion-specific models (Fig. S2).

Partial Dependence Plots (PDPs) of precipitation amount-driven subsections (Fig. 5) showed that, for most subsections, annual GPP decreased rapidly before the curves flattened at a PCI equal to 13. This suggests that years with more evenly distributed precipitation (i.e., lower PCI values) had higher GPP than years with less evenly distributed precipitation (i.e., higher PCI values), given the same amount of precipitation (Fig. 5a). Precipitation amount showed a positive control where annual GPP increased with an increase in precipitation amount before reaching a plateau at 300-700 mm/yr, depending on the subsection (Fig. 5c). In drier subsections in the CCR, GPP may decrease more than 200 g C/m^2^/yr when growing season precipitation varied from 400 to 100 mm/yr.

**Figure 5.**
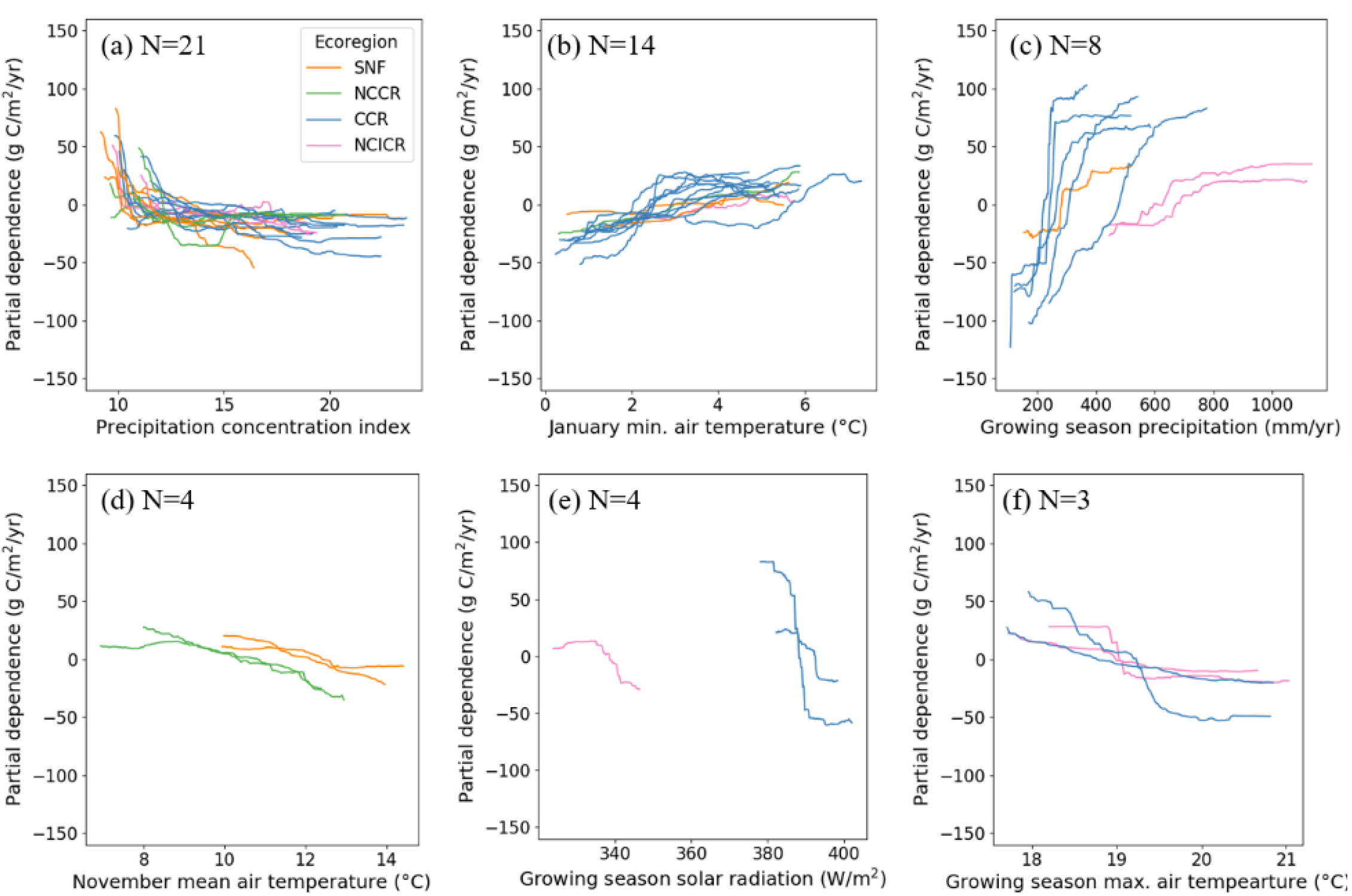
Partial dependence of annual GPP on key climate variables, including (a) precipitation concentration index (PCI), (b) January minimum air temperature, (c) growing season (October-May) precipitation amount, (d) November mean air temperature, (e) growing season solar radiation, (f) growing season mean solar radiation, and (g) growing season maximum air temperature.

Air temperatures (T_min_, T_mean_, and T_max_) showed different controls on GPP. Warmer January minimum air temperature enhanced annual GPP as it warmed from 0 to 6°C (Fig. 5b). In contrast, mean November air temperature and growing season maximum air temperature showed a negative impact on annual GPP (Figs. 5d,f). For example, as November mean air temperature increased from 7 to 14°C, GPP decreased from 10~40 g C/m^2^/yr above the long-term-mean (during 2001-2019) to 5~40 g C/m^2^/yr below the long-term-mean in subsections SNF-3, SNF-4, NCCR-3, and NCCR-4 (Fig. 5d). Maximum air temperature during the growing season showed a more substantial negative impact than mean air temperature. Annual GPP decreased more than 100 g C/m^2^/yr as growing season maximum air temperature increased from 17°C in a colder year to 21°C in a warmer year in CCR-2, CCR-3, NCICR-1, and NCICR-3 (Fig. 5f).

### 3.3 Changes in GPP under future climate projections

The ensemble mean of the four GCM projections showed an increase in total growing season (October-May) precipitation by 137 mm/yr (lower emissions) or 265 mm/yr by the end of the century under RCP 8.5, and higher monthly variation (less evenly distributed) (Fig. 6b-c & Fig. S4b-c). Precipitation projections, however, varied between GCMs. A drier climate was projected by MIROC5 versus a wetter climate by CNRM-CM5 (Fig. 7a-b). In contrast, consistent warming was projected by all GCM simulations, e.g., growing season mean air temperatures (October-May) increased by 1.7±0.7°C and 2.5±0.7°C by end of the century, under RCP 4.5 and RCP 8.5, respectively. Individual models showed a rise between 0.1°C (CNRM-CM5) and 2.8°C (HadGEM-ES) for the intermediate emissions scenario (RCP4.5) by end of the century (compared to 2001-2019), and a rise between 1.75°C (MIROC5) and 3.5°C (HadGEM-ES) for the worst-case emissions scenario (RCP8.5) in growing season mean air temperatures (October-May), with a large degree of spatial variability within the study area (Fig. 6e & Fig. S4e).

**Figure 6.**
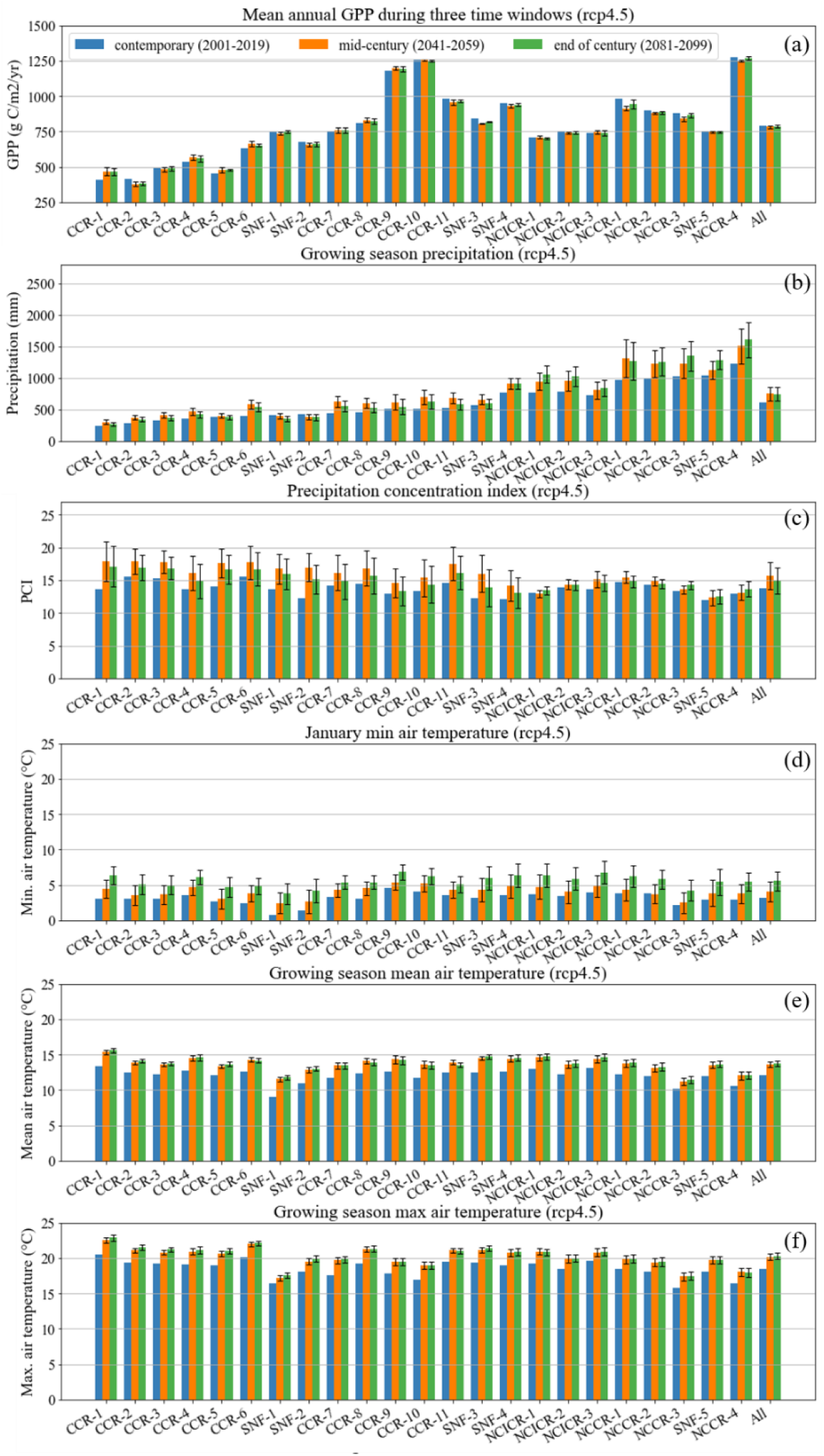
Long-term-mean (a) GPP, (b) growing season (October-May) precipitation amount, (c) growing season precipitation concentration index (PCI), (d) January minimum air temperature and growing season (e) mean and (f) maximum air temperatures during 2001-19 (contemporary), 2041-2059 (mid-century), and 2089-2099 (end of century) under the intermediate scenario (RCP4.5). Higher PCI indicates higher variability in monthly precipitation.

**Figure 7.**
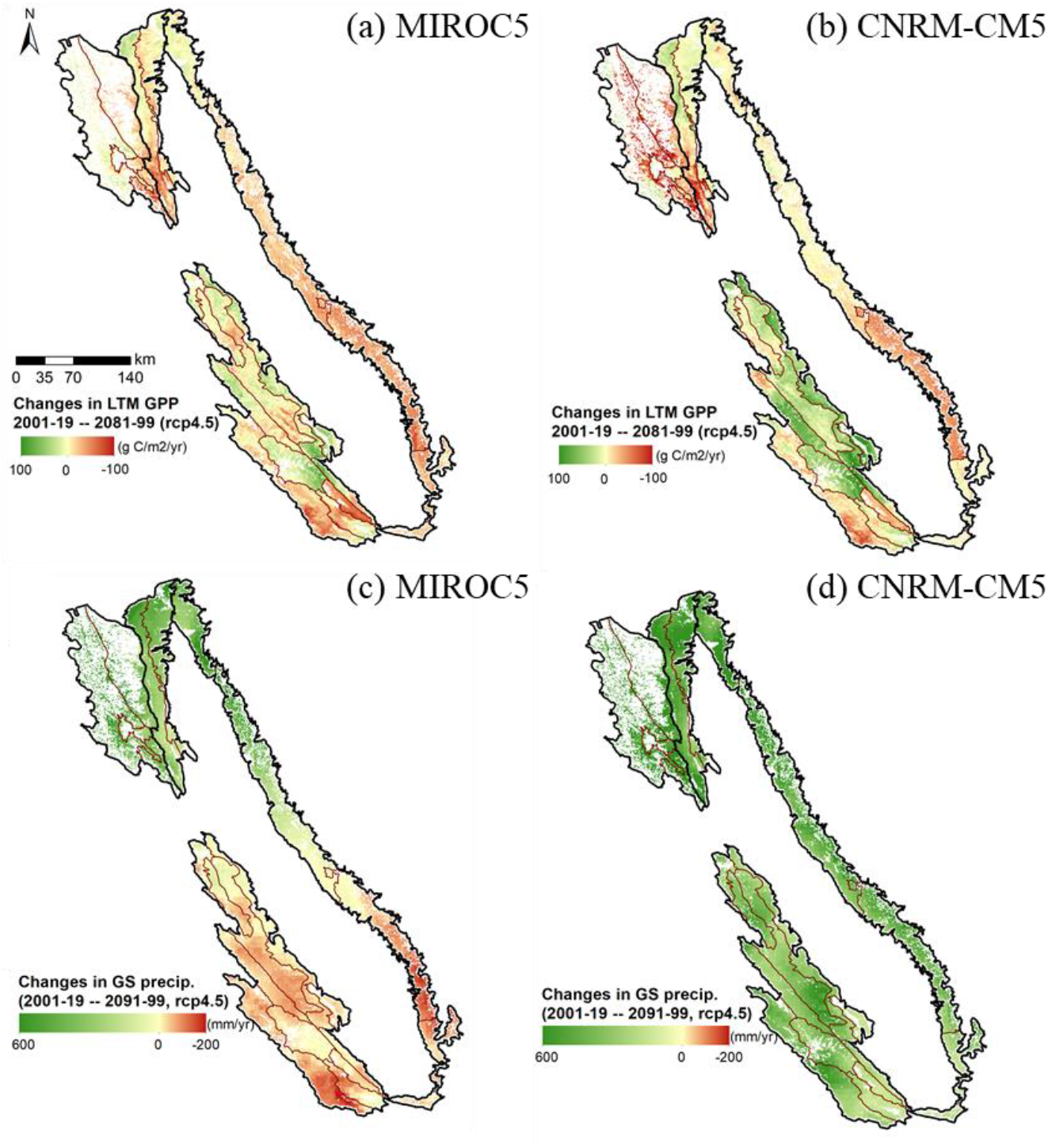
Predicted changes in GPP based on (a) MIROC5 (dry model) and (b) CNRM-CM5 (wet model), as well as the corresponding simulated changes in growing season (GS) precipitation (c-d) by the end of century under the intermediate scenario (RCP4.5).

Across the entire study area, mean annual GPP was projected to decrease from 816±271 g C/m^2^/yr to 809±260 g C/m^2^/yr by mid-century and to 812±264 g C/m^2^/yr by the end of the century (2081-2099) under RCP4.5 (Table 3). Although mean changes in GPP are small at the study area scale, these changes are not spatially uniform within the study area (Fig. 8). Under the intermediate scenario (RCP4.5), predicted GPP increased in 41% of the study area and decreased in 59% of the study area (Table 3) by the end of the century compared to the contemporary period (2001-2019). The corresponding mean increase and decrease predicted in GPP were 23±18 and −27±21 g C/m^2^/yr, respectively. Under the worst-case scenario (GCP8.5), the model predicted similar GPP changes at the study area level. However, within the study area, a larger extent of the study area is predicted to have increased GPP than under GCP4.5. The magnitude of changes in GPP is also larger under GCP8.5 (Table 3).

**Figure 8.**
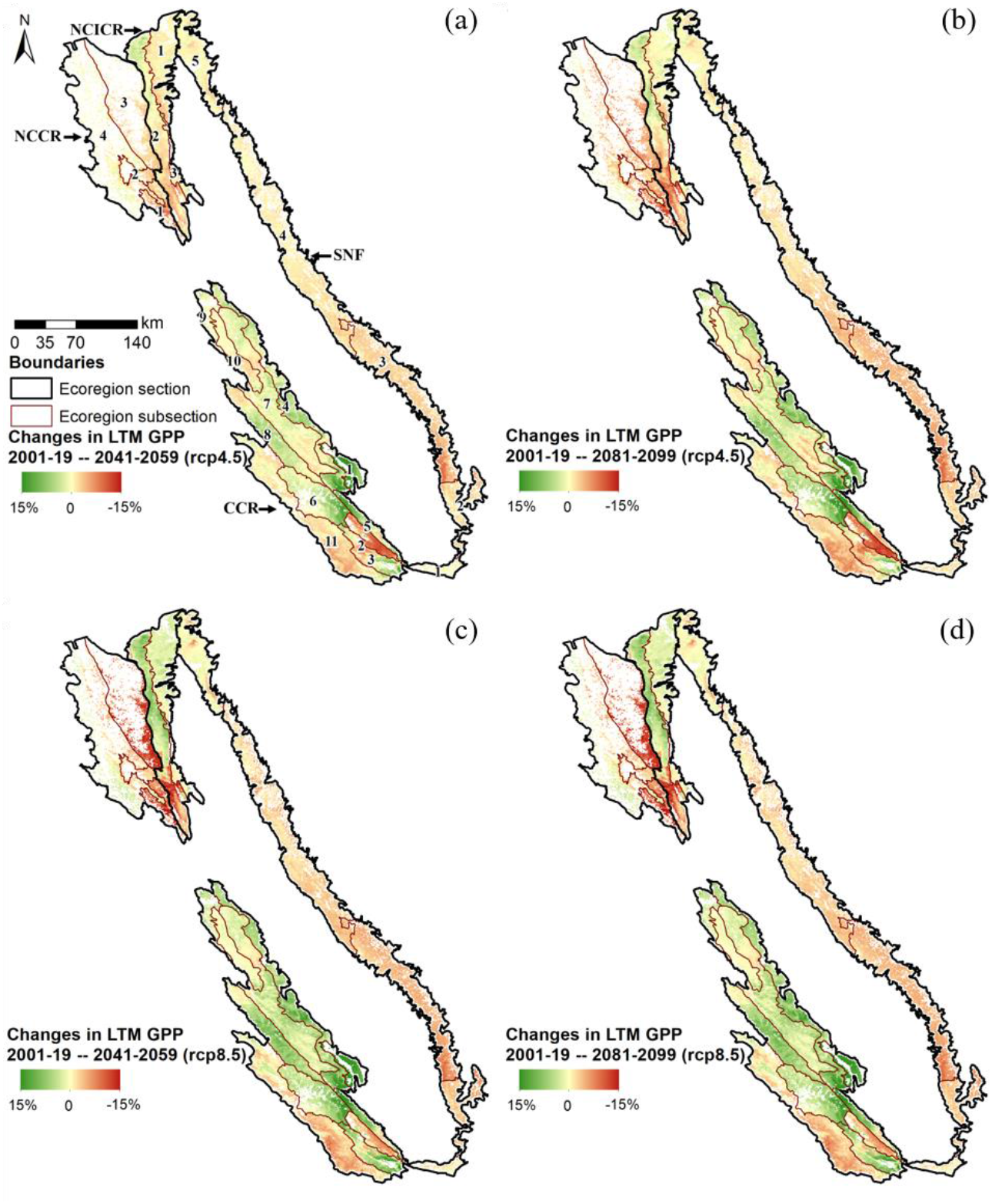
Percentage change in GPP by (a & c) mid-century and end of century (b & d) compared to the contemporary period under (a & b) intermediate (GCP4.5) and (c & d) worst-case (GCP8.5) scenarios.

**Table 3.**
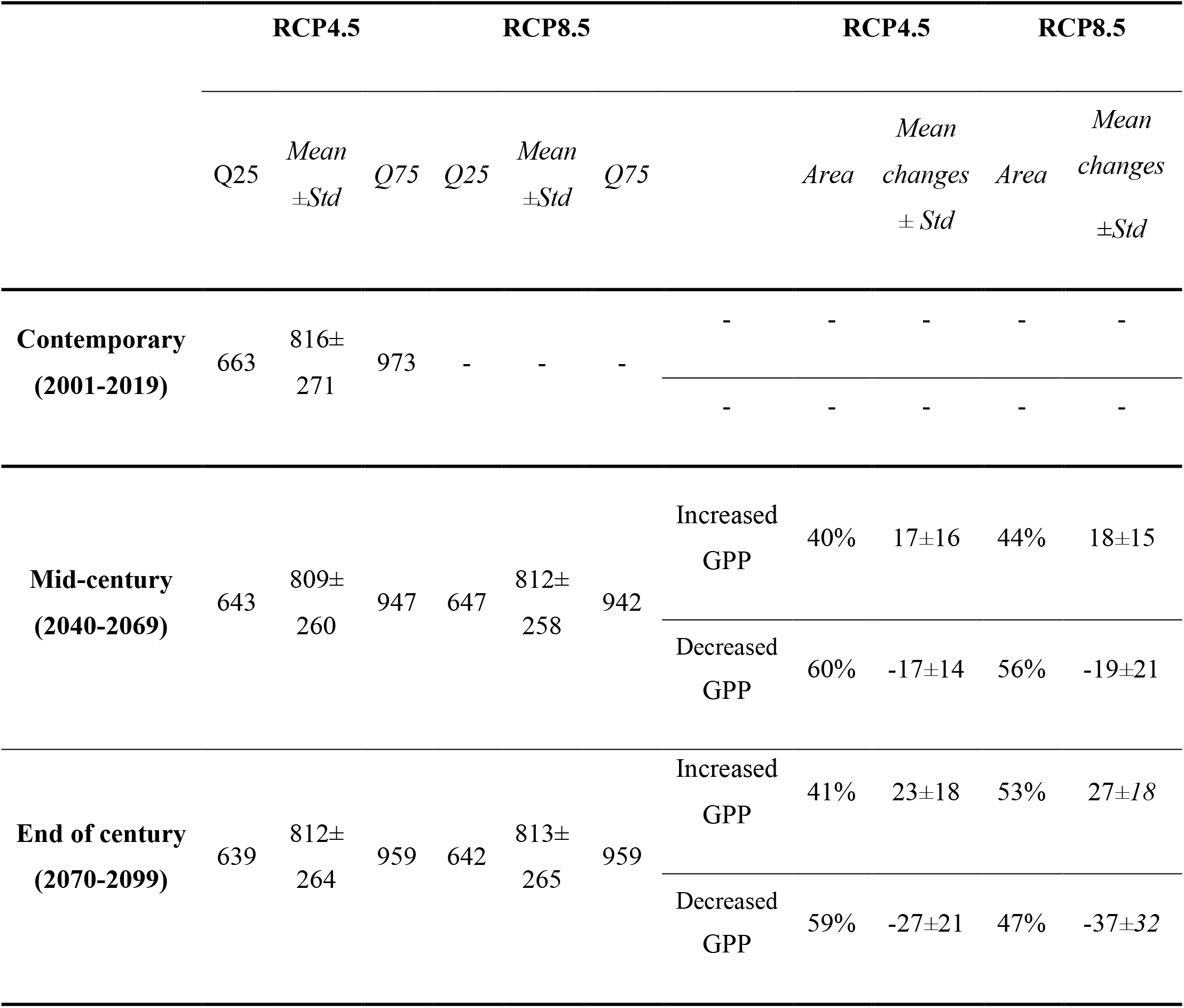
Projected GPP (g C/m^2^/yr) for the entire study area (lower and upper 25% quantiles, mean and standard deviation) during contemporary (2001-19), mid-century (2041-59), and end of century 2081-99) periods under two different emission scenarios, as compared to the contemporary time period. Areas are expressed as percentage of the entire study area.

Increases in future GPP were found mostly in drier subsections (left-hand side of Fig. 9), with a few exceptions (Fig. 9). GPP in drier subsections that received less than 513~522 mm/yr precipitation are expected to benefit from predicted increased precipitation (Fig. 6b). Increasing minimum air temperature further enhances rangeland productivity in these areas, as well as most other subsections, but reduces productivity in CCR-11, NCICR-2, SNF-5, and NCCR-1. Increased mean air temperature may bring positive or negative impacts on future GPP, depending on location. The projected increase in maximum air temperature, however, caused a decline in GPP in 4 subsections, by 45 g C/m^2^/yr in subsection CCR-2, 12 g C/m^2^/yr in subsection CCR-3, 13 g C/m^2^/yr in subsection 28 (located in NCICR), and 3 g C m^2^/yr in subsection NCICR-3. As expected, increasing PCI reduced GPP for all subsections, especially over moderate to wetter areas; GPP declined by as much as 15-30 g C/m^2^/yr in subsections CCR-11, SNF-3, and SNF-4 (more than 29% of the entire study area) where the impact of PCI was dominant. In wetter regions, reduction in GPP was further caused by an increase in growing season precipitation and warmer T_mean_.

**Figure 9.**
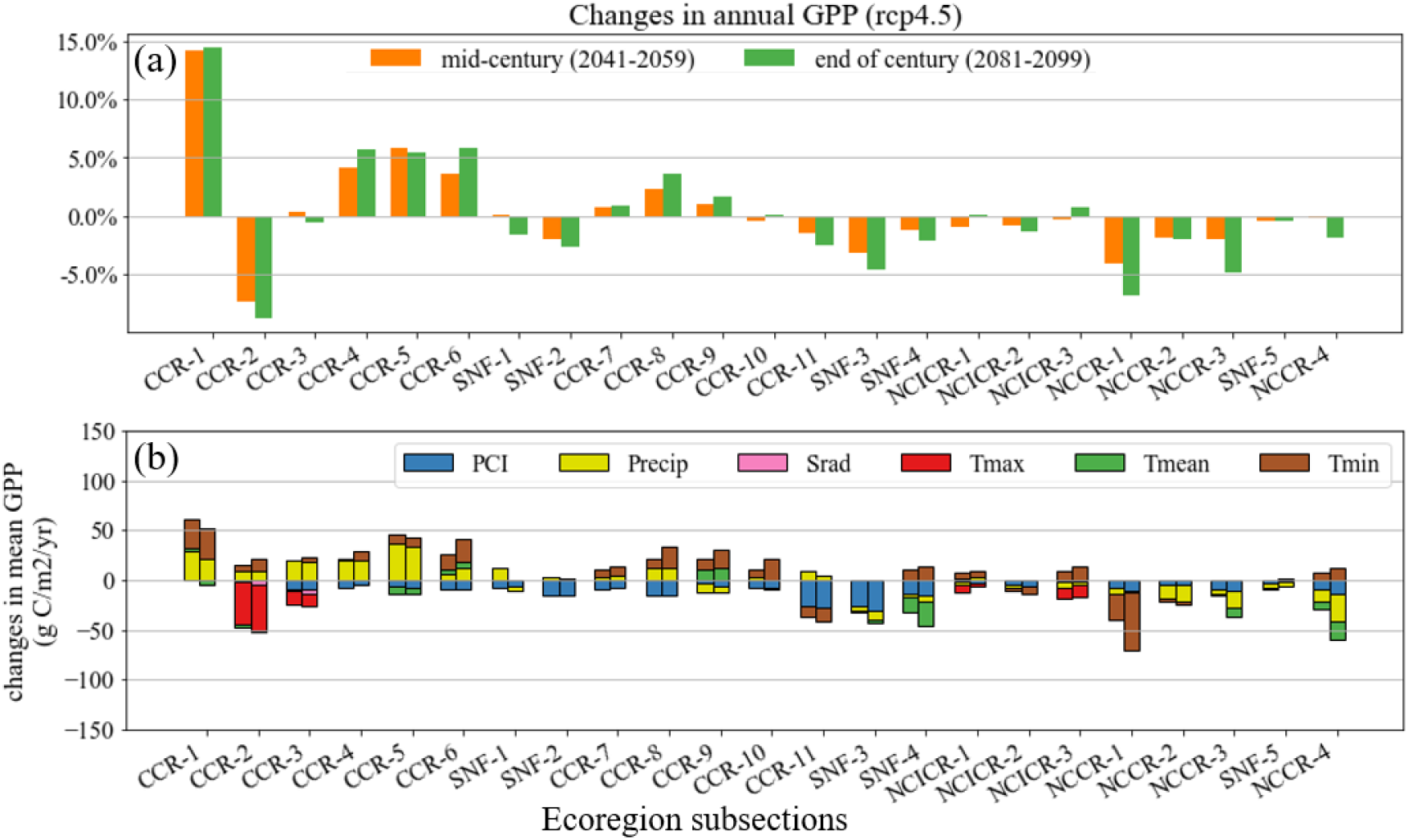
Percent change in GPP and their attribution to climatic variables for the 23 subsections under the intermediate scenario (RCP4.5). Subsections are ordered from driest (left) to wettest (right). For attribution under RCP8.5, see figure S6.

By mid-century (2041-2059), the mean annual GPP decrease across the study area was less than by the end of century under RCP4.5, but the proportions of the study area predicted with increased or decreased GPP only differed by 1% (Table 3). The actual changes in GPP values for mid-century were lower than those for the end of the century. Under the worst-case scenario (RCP8.5), predicted means for the annual GPP decrease were smaller than that of RCP4.5 by both mid-century and end of century. Also, under RCP8.5, a larger portion of the study area was predicted to experience an increase rather than a decrease in GPP compared to RCP4.5 for both time periods (Table 3). However, the magnitude of changes in GPP was larger than RCP4.5, especially by the end of the century. In summary, under the worst-case scenario, GPP is predicted to increase over a larger extent relative to the intermediate scenario. At the same time, expected changes in GPP were larger than under the intermediate scenario (Table 2 & Table 3). Attribution of these GPP changes was similar to what we discussed for RCP4.5 by end of century (Fig. S6).

Several uncertainties in GPP projections exist due to inconsistencies among GCMs, especially in precipitation simulations. Discrepancies between GPP projections by mid-century under RCP4.5 were as much as 36 g C/m^2^/yr between the MIRCO5 (dry model) and CNRM-CM5 (wet model) results (Fig. 7c-d), resulting in a large discrepancy in the GPP change attribution results (Fig. 10). For example, changes in precipitation may bring negative impacts on GPP in some drier subsections based on MIROC5, whereas in the CNRM-CM5 based results, precipitation changes are expected to have positive impacts on GPP in drier subsections. Despite the difference in GPP change attribution, the underlying relationship between GPP and precipitation is consistent because MIROC5 predicts a drier future for dry subsections while CNRM-CM5 predicts a wetter future.

**Figure 10.**
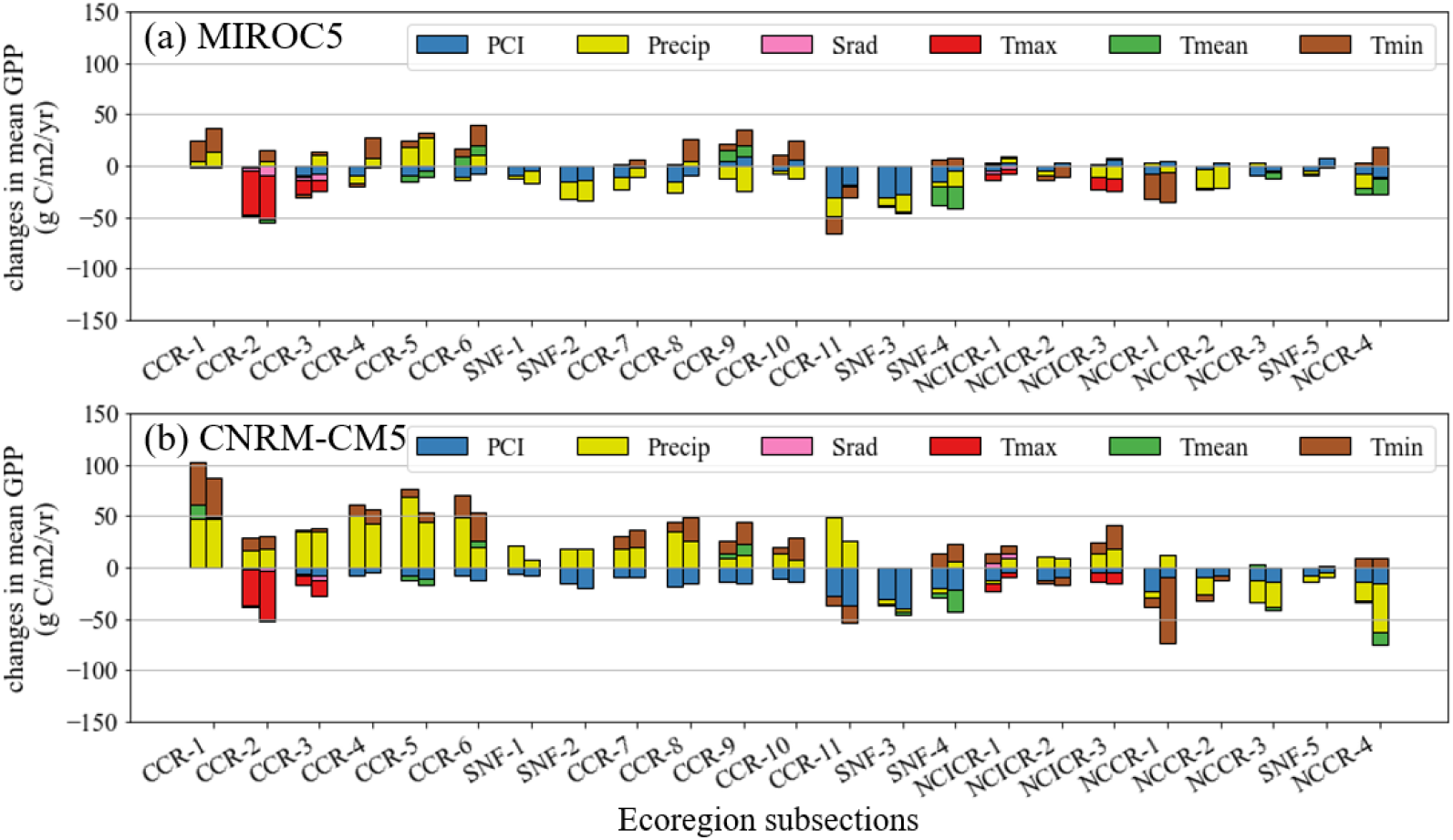
Attribution of changes in GPP for (a) MIROC5 (dry model) and (b) CNRM-CM5 (wet model) for the end of century. Subsections in the x-axis are ordered from dry (left) to wet (right) based on the contemporary period.

## 4. Discussion

### 4.1 Impacts of climate variables on rangeland GPP

This study leveraged 20 years of remote sensing records and machine learning methods to quantify and disentangle the non-linear response of annual GPP to a suite of climate variables across California’s annual rangelands. We found a high degree of spatial variability in responses across 23 ecologically distinct subsections. While interannual GPP was controlled by different climatic variables across ecoregion subsections, precipitation-based variables, in general, were major drivers of rangeland productivity responses. Moving from drier to wetter ecoregion subsections, the driving climatic variable for GPP shifted from total growing season precipitation to the relative distribution of growing season rainfall (PCI). We found when precipitation was sufficient (~300-700 mm/yr depending on ecoregion subsection), PCI was an important driver of GPP, particularly in wetter subsections. In contrast, drier ecoregion subsections were more sensitive to total growing season precipitation. Air temperatures (minimum, mean, and maximum) showed different controls on GPP. We found a positive relationship between GPP and winter minimum air temperature in all but 4 subsections (SNF-1, SNF-2, SNF-3, and NCCR-3), likely due to warmer temperatures in the early season promoting germination and early growth (Becchetti et al., 2016). In contrast, mean and maximum air temperatures showed a negative control on GPP in only nine and four subsections, respectively. Solar radiation showed a negative control on GPP, suggesting higher insolation enhanced drying processes, e.g., via evapotranspiration, in our study area. Further work is needed to better understand coupling effects of climate variables, such as the combined impacts of cold temperatures and high precipitation on GPP, as well as reasons for spatial differences in climate controls on GPP, such as why GPP is not as responsive to air temperature in some areas as it is in other areas.

### 4.2 Future changes in rangeland GPP

While our models predicted small changes in future rangeland productivity for the entire study area (Table 3), impacts at the ecoregion subsection scale were more marked and suggest rangeland productivity responses to climate change will be highly variable at the local level (Fig. 7). We found greater magnitude of GPP changes (mostly increases) in many drier subsections (Fig. 9a) in response to slightly higher expected precipitation and higher growing season minimum temperatures over time. Considering the entire study area, predicted increases in GPP from favorable future precipitation and temperature conditions in mostly drier subsections (41% of the study area) were counterbalanced by predicted decreases in GPP from unfavorable changes in temperature, rainfall distribution, and solar radiation in wetter subsections (59% of the study area)—resulting in the overall small decrease in GPP.

Several uncertainties in GPP projections exist due to inconsistencies among global climate models (GCMs), particularly for precipitation simulations (Pierce et al., 2018). Discrepancies between GPP projections by mid-century under RCP 4.5 were as much as 36 g C/m^2^/yr between the MIRCO5 (dry model) and CNRM-CM5 (wet model) results, resulting in a large discrepancy in the GPP change attribution results. For example, changes in precipitation may bring negative impacts on GPP in some drier subsections based on MIROC5, whereas in the CNRM-CM5 based results, precipitation changes are expected to have positive impacts on GPP in drier subsections. Despite the difference in GPP change attribution, the underlying relationship between GPP and precipitation is consistent because MIROC5 predicts a drier future for dry subsections, whereas CNRM-CM5 predicts a wetter future.

Climate change threatens California’s annual rangelands, which are largely rain-falldependent and therefore highly vulnerable to weather extremes (Macon et al., 2016; Roche, 2016). More extreme weather events and more frequent year-to-year weather swings in the future will lead to greater interannual rangeland productivity, potentially threatening the sustainability of forage supply and, ultimately, the resilience of ranches and rangelands to climate change. Other modeling approaches have similarly projected declines in productivity in various rangeland systems around the globe, with associated decreases in ecosystem goods and services, including livestock production (Boone et al., 2018). The high degree of spatial variability in productivity impacts projected by our models means local, climate-informed decision-making will be increasingly critical to identifying adaptation strategies for mitigating climate risks.

This work can be leveraged to create near-real time rangeland productivity tracking and forecasting tools to support adaptive decision-making across California’s extensive and highly variable rangelands. Future research and development will need to combine new data and emerging technologies with land management expertise to create locally-calibrated decision support systems. Working with land managers to co-develop decision-support tools will ensure on-the-ground relevance to sustainability and resilience at local scales and, therefore, greatly increase manager adoption. Broader benefits potentially include producing locally-trusted tools and data to help inform policy and management guidance (e.g., drought and disaster early warning systems) at regional and national scales.

## 5. Conclusion

Climate change is expected to impact forage production across California’s annual rangelands, threatening the economic viability and environmental sustainability of these highly valued systems. Our machine learning-based analysis highlighted regional differences in GPP vulnerability to climate and provided insights on the intertwining and potentially counteracting effects of seasonal temperature and precipitation regimes. This study projected the degree to which GPP may respond to changes in different climatic variables by mid-century and end of century under RCP4.5 (intermediate emissions) and RCP8.5 (worst-case emissions) scenarios using remote sensing and machine learning techniques. These results will inform rangeland management and conservation in the context of climate adaptation during an era of climate change.

## Acknowledgments

This work was supported by the Russell L. Rustici Rangeland and Cattle Research Endowment.

## Supplemental materials

**Figure S1.**
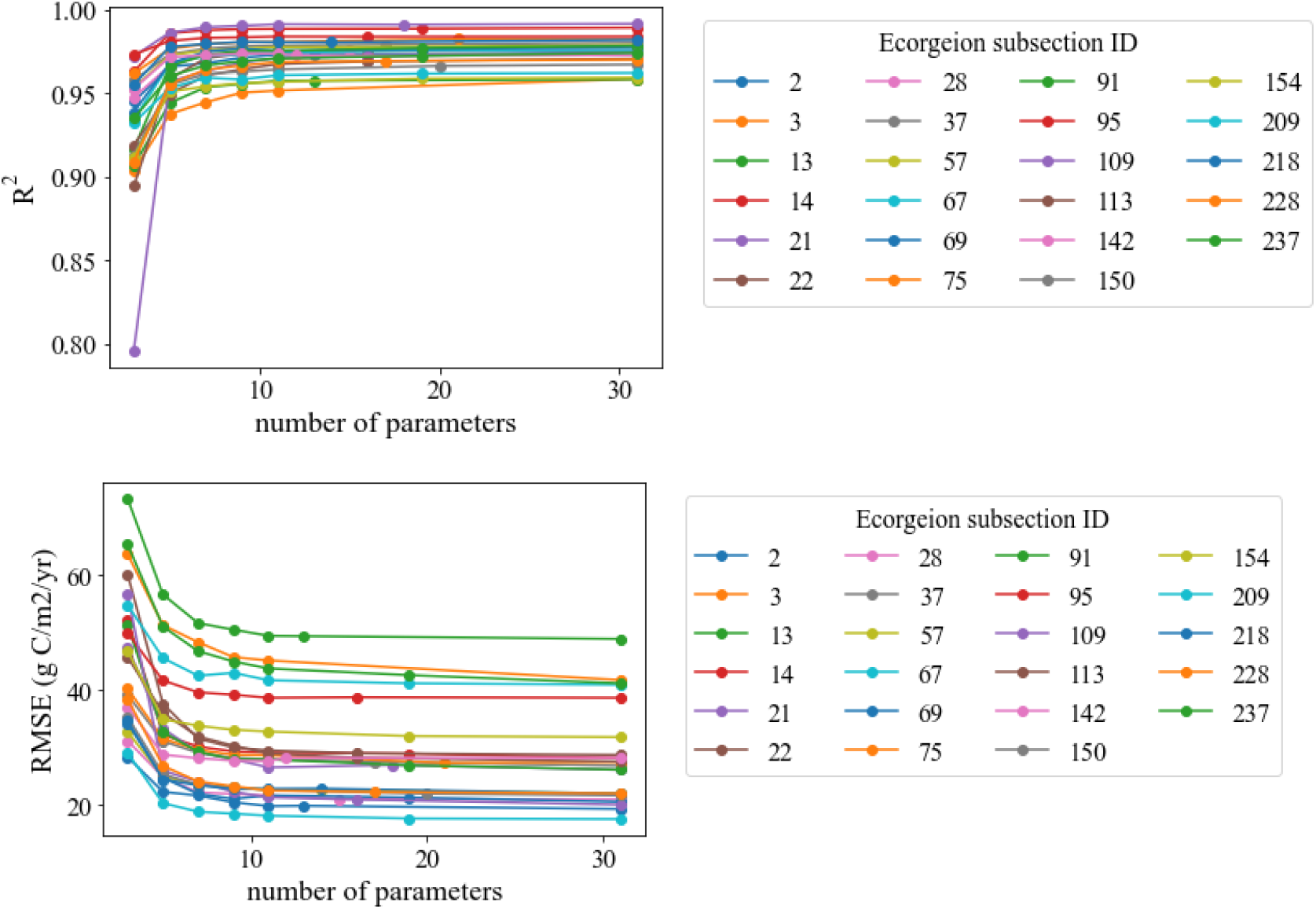
Model accuracy (R^2^ and RMSE) vs. number of features. We started with 31 features and reduced the number of features based on correlation matrix, Recursive Feature Elimination with Cross-Validation (RFECV), and finally permutation importance score.

**Figure S2.**
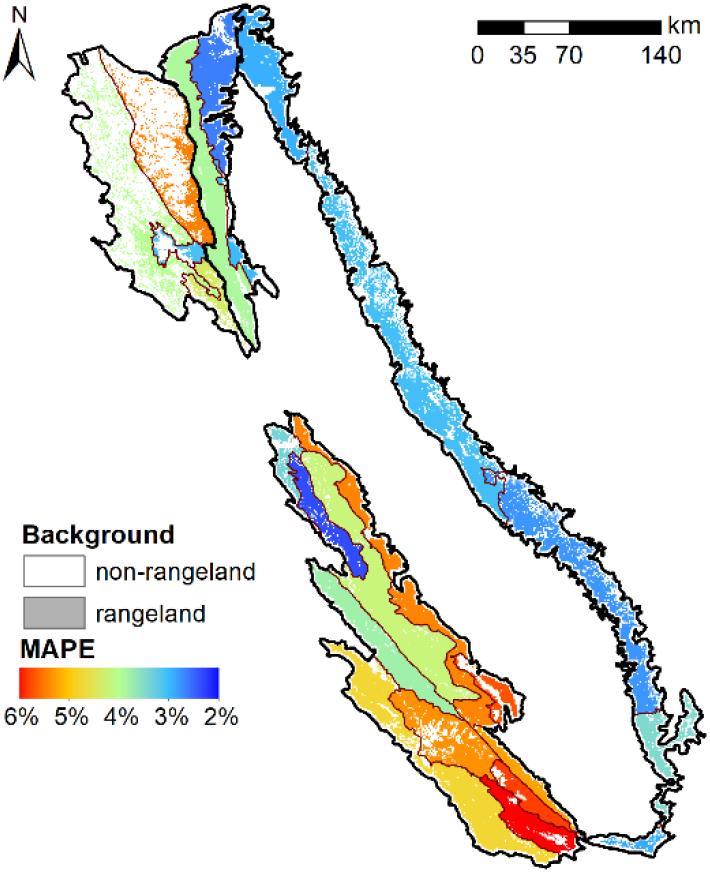
Maps of mean absolute percent error (MAPE) for each of the 23 subsection-specific models. MAPE is calculated on an independent testing dataset sampled from s subsect (25%) of each subsection.

**Figure S3.**
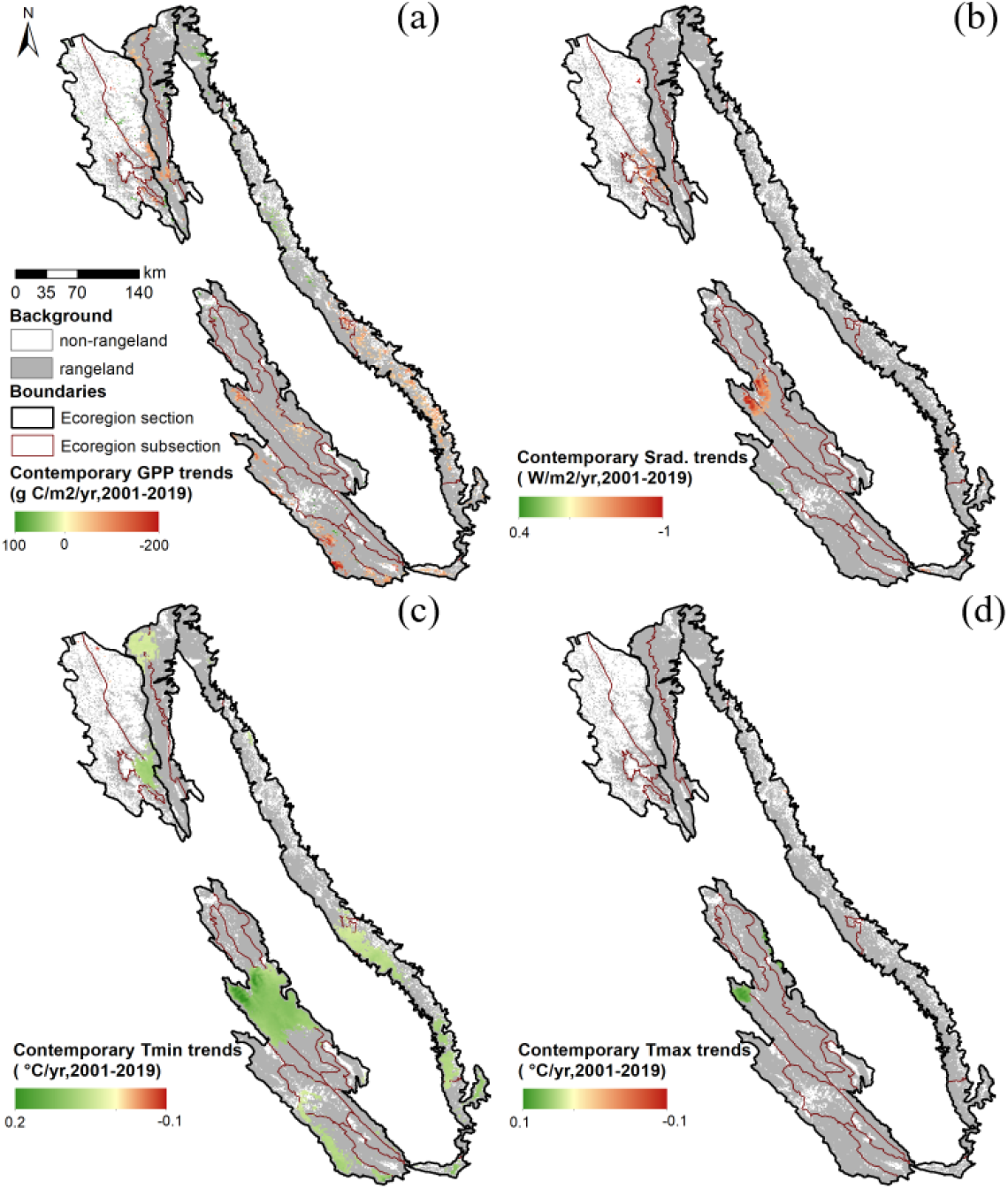
Trend maps of contemporary (a) GPP, (b) solar radiation, (c) growing season (October-May) minimum air temperature, and (d) growing season maximum air temperature.

**Figure S4.**
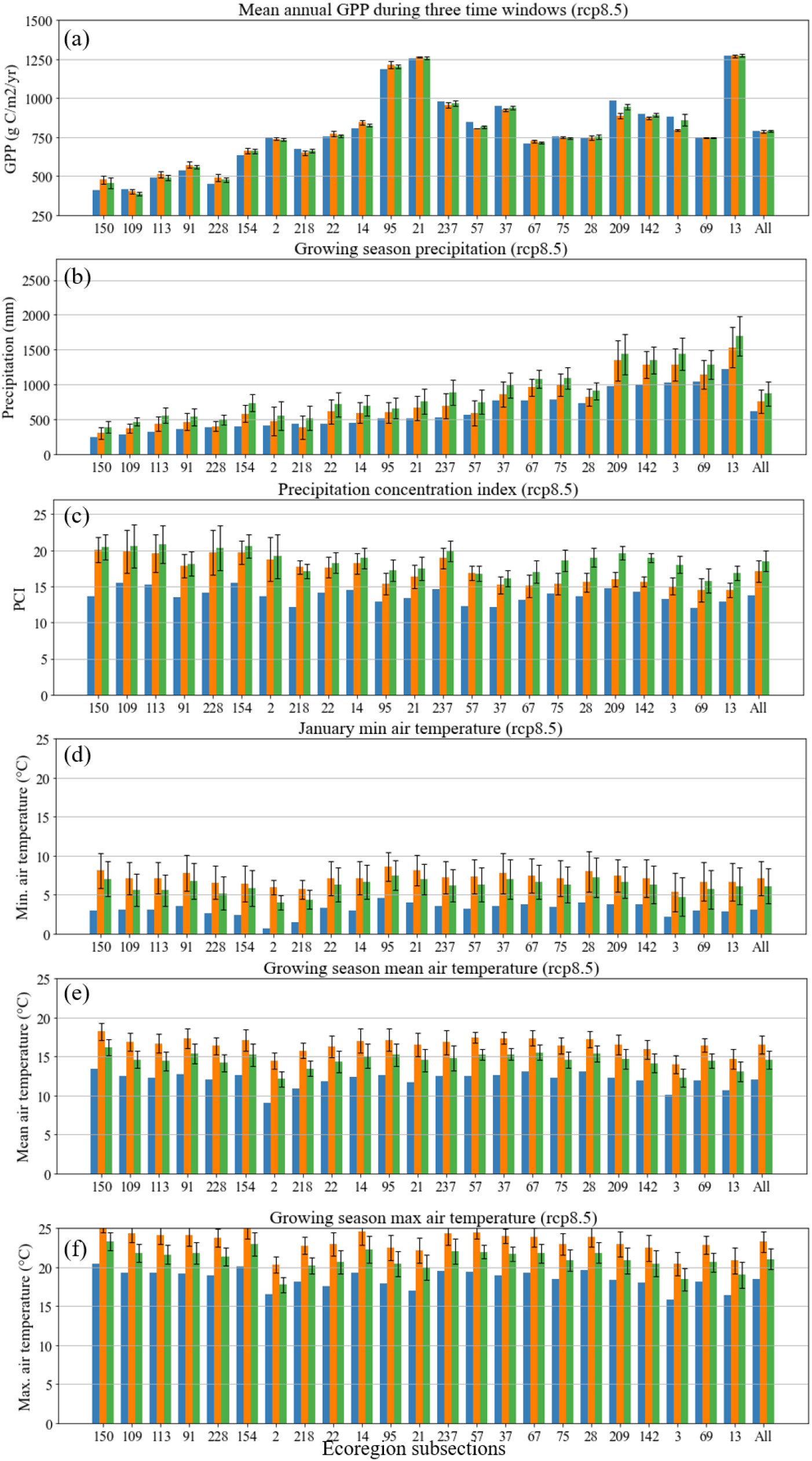
Long-term-mean (a) GPP, (b) growing season (October-May) precipitation amount, (c) growing season precipitation concentration index (PCI), (d) January minimum air temperature and growing season (e) mean and (f) maximum air temperature during 2001-19 (contemporary), 2041-2059 (mid-century), and 2089-2099 (end of century) under the worst-case scenario (RCP8.5). Higher PCI indicates higher variability in the monthly distribution of precipitation within a year. The x-axis subsections are ordered from the driest (left) to wettest (right), except for the last column presenting the entire study area.

**Figure S5.**
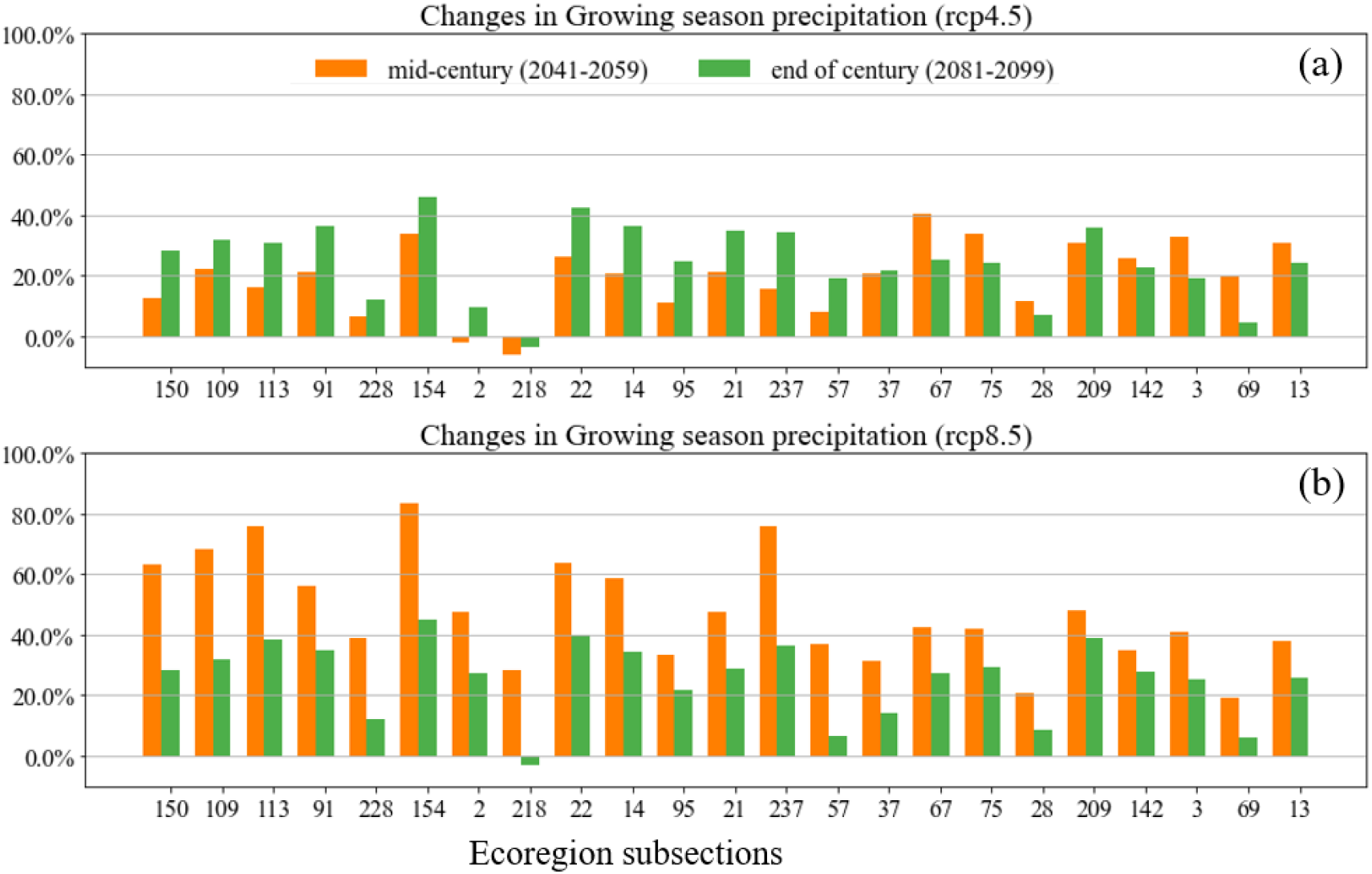
Predicted changes in growing season precipitation under (a) intermediate emissions scenario and (b) the worst-case (RCP8.5) scenario for all 23 subsections. Subsections are order from dry (left) to wet (wet) based on the contemporary period.

**Figure S6.**
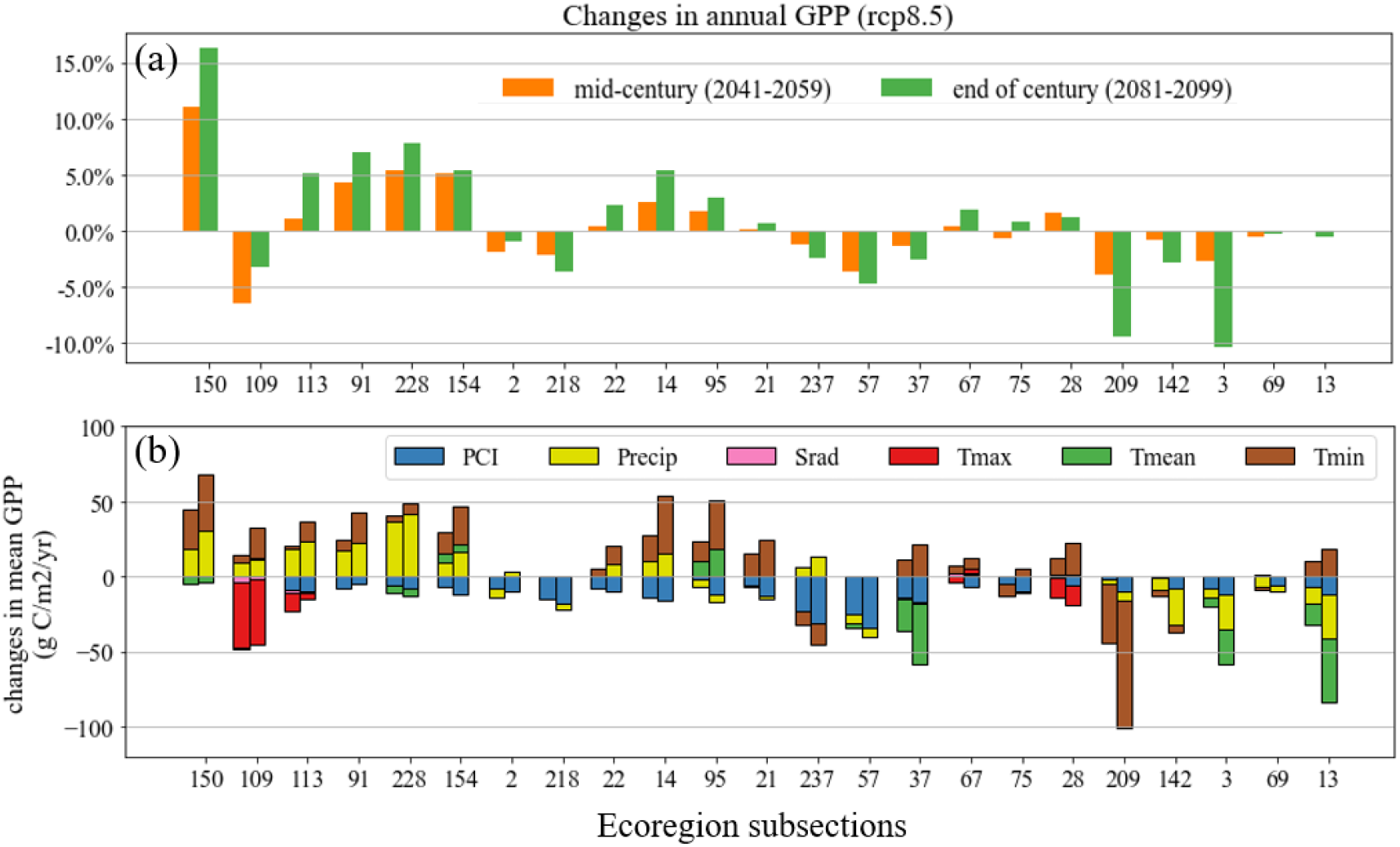
Percentage changes in GPP and their attribution to climatic variables of 23 subsections under the intermediate scenario (RCP8.5). Subsections are ordered from the driest (left) to wettest (right).

